# DHA shortage causes the early degeneration of photoreceptors and RPE in mice with peroxisomal β-oxidation deficiency

**DOI:** 10.1101/2023.08.09.552578

**Authors:** Daniëlle Swinkels, Sai Kocherlakota, Yannick Das, Adriaan D. Dane, Eric J.M. Wever, Frédéric M. Vaz, Nicolas G. Bazan, Paul P. Van Veldhoven, Myriam Baes

**Author notes:** Corresponding author: Herestraat 49 O&N2 Box 823, Leuven, 3000, Belgium.

## Abstract

**Purpose:** Patients deficient in peroxisomal β-oxidation, which is essential for the synthesis of docosahexaenoic acid (DHA, C22:6n-3) and breakdown of very long-chain polyunsaturated fatty acids (VLC-PUFAs), both important components of photoreceptor outer segments, present with retinopathy. The representative mouse model lacking the central enzyme of this pathway, multifunctional protein 2 (*Mfp2^−/−^*), also develops early onset retinal decay and cell-autonomous retinal pigment epithelium (RPE) degeneration, accompanied by reduced plasma and retinal DHA levels. In this study, we investigated whether DHA supplementation can rescue the retinal degeneration of *Mfp2^−/−^* mice.

**Methods:** *Mfp2^+/−^* breeding pairs and their offspring were fed a 0.12% DHA or control diet during gestation, lactation and until sacrifice. Offspring were analysed for retinal function via electroretinograms, for lipid composition of neural retina and plasma with lipidome analysis and gas chromatography respectively, and histologically using retinal sections and RPE flatmounts at the age of 4, 8 and 16 weeks.

**Results:** DHA supplementation to *Mfp2^−/−^* mice restored retinal DHA levels and prevented photoreceptor shortening, impaired functioning and death until 8 weeks. In addition, rescue of retinal DHA levels temporarily improved the ability of the RPE to phagocytose outer segments and delayed the RPE dedifferentiation. However, despite the initial rescue of retinal integrity, DHA supplementation could not prevent retinal degeneration at 16 weeks.

**Conclusions:** We reveal that the shortage of systemic supply of DHA is pivotal for the early retinal degeneration in *Mfp2^−/−^* mice. Furthermore, we unveil that adequate retinal DHA levels are essential for both photoreceptor and RPE homeostasis.

## Introduction

The fatty acid composition of the photoreceptor outer segment (POS) phospholipids is peculiar, as it is highly enriched in polyunsaturated fatty acids (PUFAs)^1^. The most abundant PUFA in the POS, which can amount to 50% of the phospholipid side chains, is the omega-3 fatty acid docosahexaenoic acid (DHA, C22:6n-3)^2^, but also substantial amounts of very long-chain PUFAs (VLC-PUFAs, >28 carbons) occur^3^. Both DHA and VLC-PUFAs are involved in several crucial processes in photoreceptors underscoring their importance.

Recently, the metabolism and trafficking of these lipids was extensively reviewed^4^. In short, the DHA content in the body originates from dietary sources, containing either the mature form or its precursor α-linolenic acid (ALA, 18:3n-3)^2,5^. The synthesis of DHA from ALA involves elongations, desaturations and a retroconversion, the latter executed by one cycle of peroxisomal β-oxidation, also known as the Sprecher pathway^5–7^. This process mostly takes place in the liver^8^, but can also occur in photoreceptors and the retinal pigment epithelium (RPE)^9–12^. Plasma DHA reaches the photoreceptor inner segments (PIS) after transport through the RPE, and is esterified into phospholipids, required for POS biogenesis. Alternatively, DHA can be further elongated to VLC-PUFAs, which is, amongst others, catalyzed by elongases of very long-chain fatty acids 4 (ELOVL4)^13^. During the daily phagocytosis, the lipid-rich POS are taken up by the RPE and its components are either degraded or recycled back to the PIS^14^.

Interestingly, peroxisomal β-oxidation is involved in several aspects of the metabolism of (VLC-)PUFAs. Besides its role in the synthesis of DHA, it is also involved in the breakdown of DHA and VLC-PUFAs^15^. Regarding the high abundance of peroxisomal β-oxidation enzymes in the different retinal cells^16,17^, these metabolic processes are presumed to take place in both the photoreceptors and RPE. The importance of peroxisomal β-oxidation in the retina is supported by the fact that patients with peroxisomal β-oxidation deficiency present with retinopathy^18–20^. Unfortunately, data on histological^21,22^ and lipid changes^23^ in the retina of patients with a deficiency in peroxisomes are scarce.

We recently studied the mechanism underlying the retinal degeneration in peroxisome deficiency, by analysing a mouse model lacking the central enzyme of peroxisomal β-oxidation, multifunctional protein 2 (MFP2)^24^. These *Mfp2^−/−^* mice presented with both developmental and degenerative anomalies, including 1) POS shortening at 2 weeks (w), 2) progressive photoreceptor degeneration, 3) impaired visual function (3w)^24^, and 4) RPE dedifferentiation and lysosomal dysfunction (3w) (Kocherlakota et al. manuscript in revision). Moreover, DHA-containing phospholipid species were severely depleted in both the retina and plasma of *Mfp2^−/−^* mice^24^. In contrast, mice lacking MFP2 specifically in photoreceptors (*Crx-Mfp2^−/−^* mice) displayed intact retinal DHA levels and photoreceptor development^25^. These findings hinted at the importance of the systemic supply of DHA for retinal DHA levels and integrity in *Mfp2^−/−^*mice.

Therefore, we aimed to restore the systemic supply of DHA in *Mfp2^−/−^* mice, by supplementing a 0.12% (w/w) DHA diet. This treatment replenished retinal DHA levels in the *Mfp2^−/−^*retina and prevented photoreceptor shortening, death and impaired functioning at early ages. Surprisingly, DHA supplementation also improved RPE morphology and functioning. However, improvements in photoreceptor and RPE homeostasis were only temporary, regardless of continuous DHA supplementation, alerting that DHA supplementation alone will not be sufficient to prevent vision loss in MFP2-deficient patients. Altogether, these data provide a better understanding of the synthesis, trafficking and function of DHA in the retina and shed new light on the pathogenesis of the retinopathy in peroxisome-deficient patients.

## Materials and methods

### Mouse handling

Global *Mfp2* knockout mice (Swiss background) were generated by breeding heterozygous mice^26^. As no retinal differences were observed between *Mfp2^+/+^* and *Mfp2^+/−^*mice, both served as control. Genotyping for *Mfp2^−/−^* and the spontaneously occurring *rd1* mutation (*Pde6* gene) was performed as described (Supplementary Table 1)^24,27^.

Animals were bred in the conventional animal housing facility of the KU Leuven and were kept on a 13–11h light and dark cycle. Experiments conformed to the ARVO statement for the Use of Animals in Ophthalmic and Vision Research and were approved by the Research Ethical Committee of the KU Leuven (P166/2017 and P129/2022). Mice were sedated with a mix of Nimatek (75 mg/kg) and Domitor (1 mg/kg). To collect plasma, the eye was removed, followed by collection of blood in heparin pre-treated tubes and centrifugation at 1200xg for 10min at 4°C. The neural retina and RPE were isolated as previously explained, 7h after light onset^24,28^.

### Dietary intervention

Swiss *Mfp2^+/−^* pregnant female mice and their offspring were fed a 0.12% DHA or control diet (Supplementary Figure 1). The dietary DHA supplements (TG-form) (MEG3®) were kindly provided by DSM Nutritional Products (Basel, Switzerland) and incorporated into the chow at Ssniff (Soest, Germany). The concentration and composition of the diet was based on Connor et al.^29^.

To determine the fatty acid composition of the diets, they were extracted via the Bligh-Dyer method and analyzed by gas chromatography (GC)^30,31^. To this end, the diet was finely ground, dissolved in methanol/chloroform (2:1 v/v) (1 g diet/3 ml solvent) and the liquid phase was collected. Next, 1M NaCl/methanol (7:3 v/v) was added, the upper phase was removed, while the lower phase was dried in the Genevac EZ-2 (SYSMEX, Belgium) and send for GC analysis.

The fatty acids in the control diet mainly consisted of palmitic acid (C16:0), oleic acid (C18:1n-9) and linoleic acid (C18:2n-6) (Supplementary Table 2), whereas DHA (C22:6n-3) could not be detected. The DHA diet comprised the same main lipid species as the control diet, but was partly substituted with 1.2% (w/w of total fatty acids) DHA. As the diets contained a total of 10% fat, the DHA diet consisted of 0.12% (w/w) DHA.

### Electroretinogram

To measure visual functionality, electroretinograms (ERGs) were performed using the Celeris system (Diagnosys), in collaboration with the laboratory of Animal Physiology and Neurobiology, KU Leuven^25^.

### Lipid measurements

To determine the total fatty acids, GC analysis was performed by the Amsterdam UMC as previously defined^32^. After correction using an internal standard, fatty acid levels were expressed as percentage of total fatty acids (diet) or µmol/L extract (plasma).

Lipidome analysis on *Mfp2^−/−^* neural retina samples was performed by the Core Facility Metabolomics of the Amsterdam UMC, as described previously^25,33^. Of note, comparisons could only be performed between different groups (e.g. WT vs. *Mfp2^−/−^*) within the same lipid classes. Data are presented as fold change compared to WT levels on control diet.

### Histological assessments

Enucleated eyes were fixed overnight at 4°C in new Davidson’s fixative (NDF) (22.2% (v/v) formaldehyde 10%, 32% (v/v) alcohol, 11.1% (v/v) glacial acetic acid), after which they were cut in 7 µm thick transverse retinal sections.

Gross morphology was assessed with haematoxylin–eosin (H&E) staining^25^. Based on Lobanova et al.^34^, the number of photoreceptor nuclei were counted over a distance of 100 µm at 6 different regions on both sides (nasal and temporal) of the optic nerve head, after which the number of nuclei were summed up. Images were acquired with an inverted IX-81 microscope (Olympus, 20× objective).

To measure the length of the photoreceptor layer (PR), POS and PIS, phase contrast microscopy (PCM) was performed as previously explained^25^. Images were acquired with a Leica DMI6000B microscope, using phase-contrast settings (63× objective).

Immunohistochemical (IHC) analysis was performed on NDF sections and RPE flatmounts as described^25^. Primary antibodies are listed in Supplementary Table 3. Images were acquired with a Leica SP8× confocal microscope. To calculate the number of rhodopsin-positive POS over a distance of 100 µm, per mouse one image was taken on either side of the optic nerve (100x objective), after which POS were counted using ImageJ (NIH) and the average was used for analysis^35^. Of note, the rhodopsin B630 variant was used, which recognizes the N-terminus of rhodopsin that remains intact until fusion with and degradation in lysosomes^36^.

### Immunoblotting

Neural retinas were homogenized as described before^25^. RPEs were homogenized using a pestle homogenizer for 30 seconds, after which the sclera was discarded. Next, immunoblotting was performed using the described protocol^25^, with the exception of the blocking buffer (5% (w/v) bovine serum albumin in 0.1% (v/v) tween20). Primary antibodies are listed in Supplementary Table 3. Images were processed with the Image Lab software (Bio-Rad). Vinculin served as loading control.

### RNA isolation and quantitative RT-PCR

The neural retina was homogenized in trizol (ThermoFisher Scientific) with a sonicator. The RPE was homogenized in lysis buffer (PureLink RNA Mini kit, Thermo Scientific) containing 2-mercaptoethanol using a pestle homogenizer for 30 seconds, after which supernatants was collected. Next, RNA was extracted, converted to cDNA and RT-qPCR was performed as previously described^24^. To calculate the relative expression to a reference gene (*Actb*), the 2^−ΔΔCT^-method was used. Primers are listed in Supplementary Table 4.

### Statistics and Reproducibility

Statistical analysis was performed using the GraphPad Prism software (version 9.3). Grubbs test was executed on every data set to identify possible outliers, Shapiro–Wilk test was used to assess normal distribution and the F-test was performed to test equality of the variances. Statistical significance was set at p<0.05 and data are presented as mean ± SD. Two-way ANOVA was used to analyze ERG responses, while one-way ANOVA with multiple comparison was performed for the other tests.

## Results

### Supplementing a DHA diet to *Mfp2^−/−^* mice normalizes plasma and neural retina DHA levels

Our previous findings of lowered plasma and retinal DHA levels in *Mfp2^−/−^* mice^24^, urged to investigate whether levels could be restored by supplementing DHA to the diet. To this end, a 0.12% (w/w) DHA or control diet was fed to *Mfp2^+/−^* pregnant female mice and their offspring until sacrifice (Supplementary Figure 1). Of note, previous studies describing the retinal phenotype of *Mfp2^−/−^* mice were performed on C57Bl6 mice^24^, while these supplementation studies were done on Swiss *Mfp2^−/−^* mice because of the larger litters.

Firstly, we assessed DHA levels in plasma of 4-week-old mice by GC analysis. As expected, DHA supplementation to *Mfp2^−/−^* mice increased plasma DHA, which even rose above levels in WT mice on control diet (4-fold) (Figure 1a). Notably, plasma DHA reached almost 2-fold higher levels in WT mice on DHA diet than in *Mfp2^−/−^* mice on the same diet.

**Figure 1.**
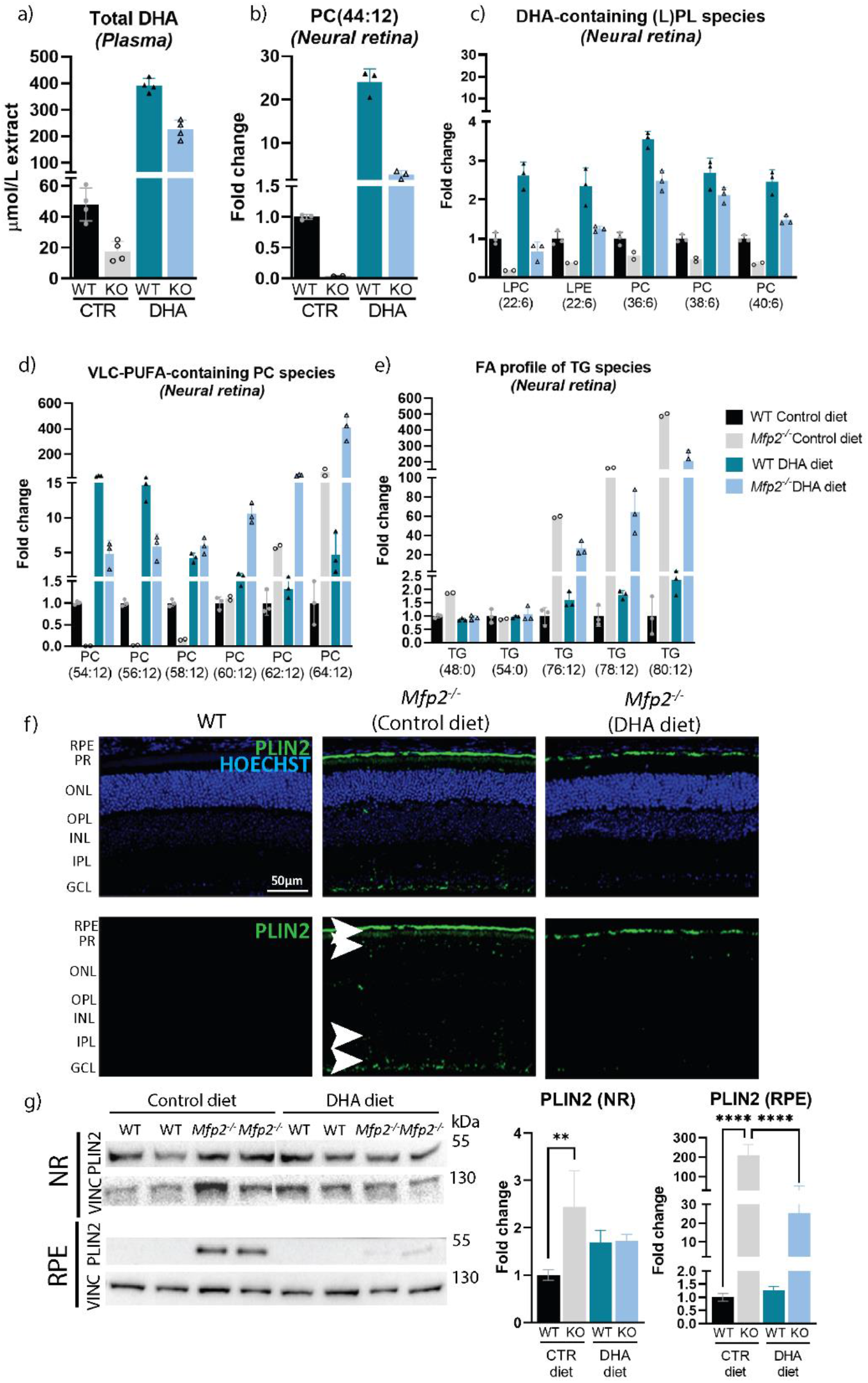
Altered lipid profile in DHA supplemented Mfp2^−/−^ mice (4w). **(a)** GC analysis to determine DHA levels in plasma. **(b)** Lipidome analysis represented as fold change for (lyso-)phospholipids ((L)PL) most likely containing two DHA moieties**, (c)** or one DHA**. (d)** VLC-PUFA-containing phospholipid species and **(e)** triglycerides. N=2-4/group. No statistical test was performed for the lipidome data, but individual data points are shown. **(f)** IHC staining for the lipid droplet marker PLIN2 (green) on 4-week-old mice. The lower panel serves to visualize the lipid droplets in the neural retina. **(g)** Immunoblotting and quantification of PLIN2 levels in 4-week-old neural retina and RPE. Vinculin was used as loading control. N=4/group. Statistical difference for immunoblotting based on multiple one-way ANOVA. Error bars indicate SD. RPE—retinal pigment epithelium; PR—photoreceptor; ONL—outer nuclear layer; OPL—outer plexiform layer; INL—inner nuclear layer; IPL— inner plexiform layer; GCL—ganglion cell layer; PC—phosphatidylcholine; PL—phospholipids; TG— triglycerides; DHA—docosahexaenoic acid; AA—arachidonic acid; VLC-PUFA—very long chain polyunsaturated fatty acid; PLIN2—perilipin 2; NR—neural retina; VINC—vinculin; ** p < 0.01; **** p < 0.0001.

Subsequently, we evaluated the effect of DHA supplementation via an extensive lipidome analysis on 4-week-old neural retinas, as DHA is primarily localized to the POS. Examination of glycerophospholipid species such as PC(44:12), which is generally accepted to contain two DHA moieties, revealed a 2.7-fold increase in the neural retina of *Mfp2^−/−^* mice on DHA diet compared to levels of WT mice on control diet (Figure 1b). Furthermore, other phospholipid species most likely containing DHA (based on the presence of 6 double bonds), increased similarly (Figure 1c). Due to technical limitations during the COVID19 pandemic, only 2 to 3 replicates could be used for lipidomic analysis, restricting the possibility of performing statistical tests. Nevertheless, the data appear to be representative, as i) there is low variation between the replicates and ii) the lipid changes in the *Mfp2^−/−^* retinas on control diet are in coherence with previously reported changes in C57Bl6 *Mfp2^−/−^* mice^24^.

The PC phospholipids were further analyzed with regard to incorporation of VLC-PUFAs, as those were previously reported to show a peculiar profile in the neural retina of C57Bl6 *Mfp2^−/−^* mice^24^. PC species, most likely composed of one DHA and one VLC-PUFA moiety, up to a length of 36 carbons (e.g. PC(58:12)), were almost absent in the neural retina of *Mfp2^−/−^* mice on control diet (80-90% reduced), but reached levels above normal in the DHA supplemented *Mfp2^−/−^* mice (5-10-fold increase), compared to WT mice on control diet (Figure 1d). This implies that the reduced VLC-PUFA levels (≤C36) in the *Mfp2^−/−^*retina were due to the lack of the precursor DHA. On the contrary, PC-species containing longer VLC-PUFAs (≥C38 (e.g. PC(60:12)) accumulated in *Mfp2^−/−^* mice on either diet, but this was more pronounced in the DHA supplemented *Mfp2^−/−^* mice, likely due to uncontrolled PUFA elongation.

Considering that VLC-PUFAs can also be incorporated into other lipid species in the neural retina^25^, the PUFA composition of triglycerides (TG) was examined. TG species presumably containing one common saturated fatty acid (e.g. C16:0 or C18:0), one DHA moiety (C22:6), and one VLC-PUFA elongated from DHA (e.g. C38:6, C40:6 or C42:6), accumulated to a lesser extent (2-3-fold) in the neural retina of *Mfp2^−/−^* mice on DHA diet compared to *Mfp2^−/−^* mice on control diet (Figure 1e). Nevertheless, levels of these TG species were still increased by 20-200-fold compared to WT mice on control diet. On the other hand, levels of TG species containing saturated fatty acids, such as 3 times C16:0 (i.e. TG(48:0)) or 3 times C18:0 (TG(54:0)) were unaltered.

Although VLC-PUFAs are enriched in the photoreceptors, it should be kept in mind that the levels of PUFAs (≥ C24) are several orders of magnitude lower than long-chain saturated or monounsaturated fatty acids^37^. However, our analysis does not allow to compare the absolute concentrations of different lipid species given the semiquantitative nature of the measurement. Therefore, we evaluated if the altered composition of the neutral lipids in the neural retina impacted on the abundance of lipid droplets, by performing an IHC staining for the lipid droplet marker perilipin-2 (PLIN2) (4w). While no lipid droplets were detected in WT mice on both diets, PLIN2 staining was clearly present in the photoreceptors, inner plexiform and ganglion cell layer of the neural retina of *Mfp2^−/−^* mice on control diet (Figure 1f). Interestingly, DHA supplementation reduced the neutral lipid accumulation in the entire neural retina of *Mfp2^−/−^* mice (Figure 1f), compared to *Mfp2^−/−^* mice on control diet. This was quantified using immunoblotting for PLIN2, revealing normalization of PLIN2 in the neural retina of DHA supplemented *Mfp2^−/−^* mice compared to WT mice (Figure 1g).

The PLIN2 staining also showed extensive lipid droplet accumulation in the *Mfp2^−/−^* RPE, similar to previous findings in C57Bl6 mice (Kocherlakota et al. manuscript in revision). Remarkably, similar to the neural retina, DHA supplementation considerably reduced the number of lipid droplets in the RPE, shown by PLIN2 IHC and immunoblotting (90% reduction) (Figure 1f-g).

Overall, the lipid analyses revealed that supplementing a 0.12% (w/w) DHA diet to *Mfp2^−/−^* mice enhanced the systemic supply of DHA to the retina, thereby restoring levels of DHA-containing phospholipid species in the neural retina. This also affected the levels and distribution of the elongation products.

### Normalizing retinal DHA levels improves photoreceptor development and delays retinal degeneration in *Mfp2^−/−^* mice

*Mfp2^−/−^* mice in the C57Bl6 background presented with POS shortening already at the age of 2w and loss of photoreceptors at 8w^24^. To explore whether restoring DHA levels in the *Mfp2^−/−^* retina could prevent these retinal abnormalities, H&E stainings (Figure 2) and morphometric analysis (Supplementary Figure 2) on retinas of 4-, 8- and 16-week-old DHA supplemented *Mfp2^−/−^* mice were performed. Interestingly, while *Mfp2^−/−^* mice on control diet presented with 30% shorter POS at 4w, this was not observed in *Mfp2^−/−^* mice on DHA diet (Figure 2a and Supplementary Figure 2a). Even more striking were the findings at 8w. The severe shortening (60%) and loss of photoreceptors (50%) in *Mfp2^−/−^* mice on control diet was prevented by supplementation with DHA (Figure 2b and Supplementary Figure 2b). However, despite the initial rescue of retinal integrity, DHA supplementation could not prevent a substantial retinal degeneration at 16w. Nevertheless, POS length and photoreceptor survival were still better preserved compared to *Mfp2^−/−^* mice on control diet at the same age (Figures 2c and Supplementary Figure 2c). Of note, no differences in retinal morphology were observed between WT mice on either diet (Figure 2a-c, quantifications). Taken together, these results indicate that impaired systemic delivery of DHA to the neural retina caused the early onset photoreceptor developmental and degeneration problems in *Mfp2^−/−^* mice. Nevertheless, other mechanisms are also at play, as the retina deteriorated at 16w, despite continuous DHA supplementation.

**Figure 2.**
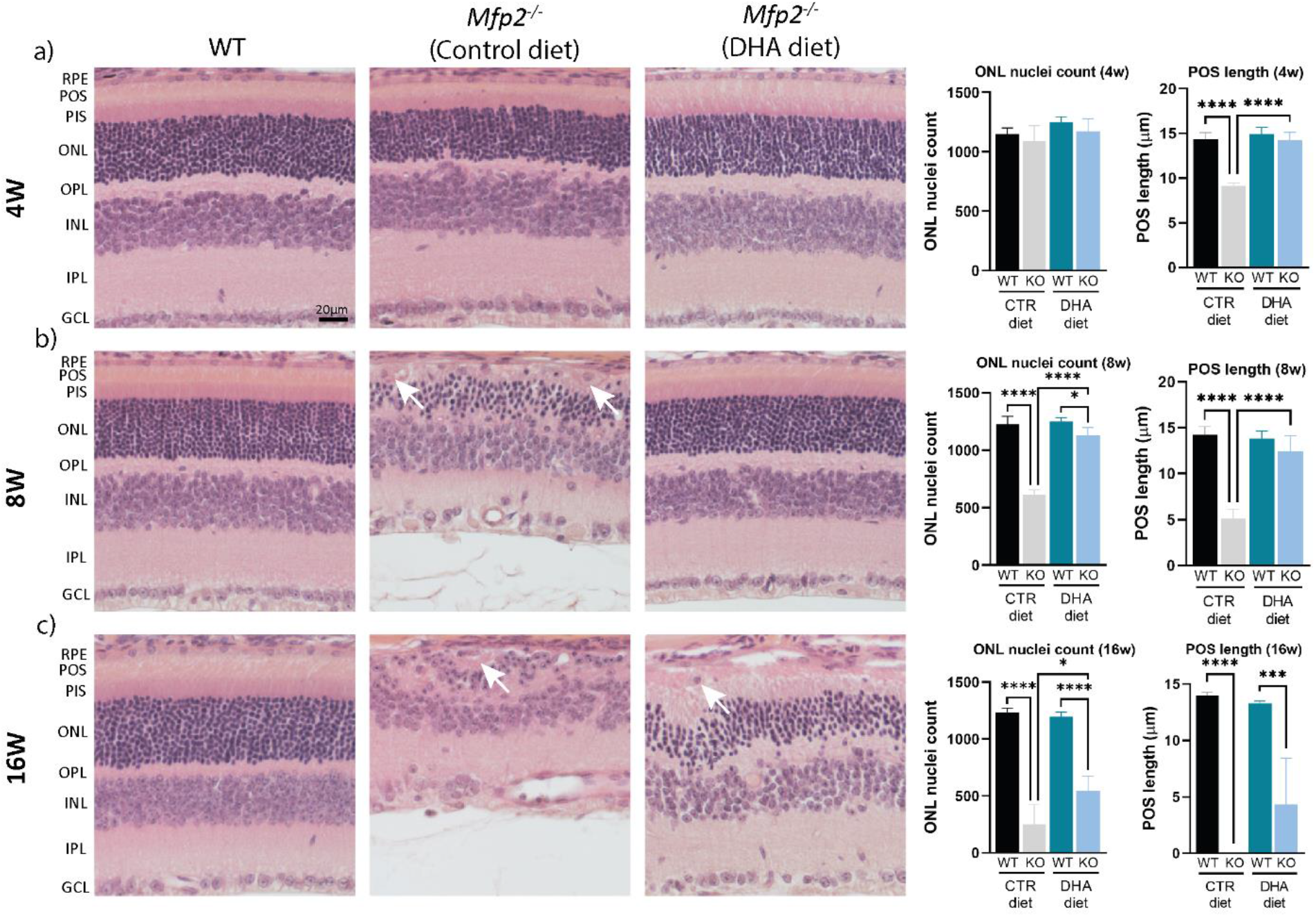
DHA supplementation to Mfp2^−/−^ mice delays retinal degeneration. H&E stainings on 4-**(a)**, 8-**(b),** and 16-week-old **(c)** mice. The right panel represent quantification of the ONL nuclei count and POS length per respective age. White arrows indicate dedifferentiated RPE cells. N=4-5/group. Statistical difference based on multiple one-way ANOVA. Error bars indicate SD. RPE—retinal pigment epithelium; POS—photoreceptor outer segments; PIS—photoreceptor inner segments; ONL—outer nuclear layer; OPL—outer plexiform layer; INL— inner nuclear layer; IPL—inner plexiform layer; GCL—ganglion cell layer; CTR—control; * p < 0.05; *** p < 0.001; **** p < 0.0001.

### Supplementation of DHA improves *Mfp2^−/−^* visual function

Next, we evaluated if restored retinal DHA levels and photoreceptor integrity in juvenile *Mfp2^−/−^* mice impacted on the function of rods, the dominant photoreceptors in the mouse retina, by measuring ERGs. Scotopic (i.e. dark-adapted) a-wave responses, which represent the activity of rod photoreceptors, of 4-week-old *Mfp2^−/−^*mice on control diet were reduced (Figure 3a), similar to previous observations in knockouts in the C57Bl6 background^24^. DHA supplementation normalized the rod photoreceptor response at the highest intensity in 4-week-old *Mfp2^−/−^* mice, which is in accordance with the intact morphology of these neurons at this age (Figure 3a). However, already at 8w the a-wave response deteriorated, despite normal retinal morphology at this age, although responses were still ±4-fold higher than in *Mfp2^−/−^* mice on control diet (Figure 3b). The scotopic b-wave response, which represents the activity of rod interneurons, was almost absent in 4- and 8-week-old *Mfp2^−/−^* mice on control diet (Figure 3a-b). The DHA diet improved the interneuron response, but it could not normalize them to wildtype levels. These findings are in line with our previous observations in *Crx-Mfp2^−/−^*mice, i.e. normal photoreceptor responses, but impaired interneuron responses^25^.

**Figure 3.**
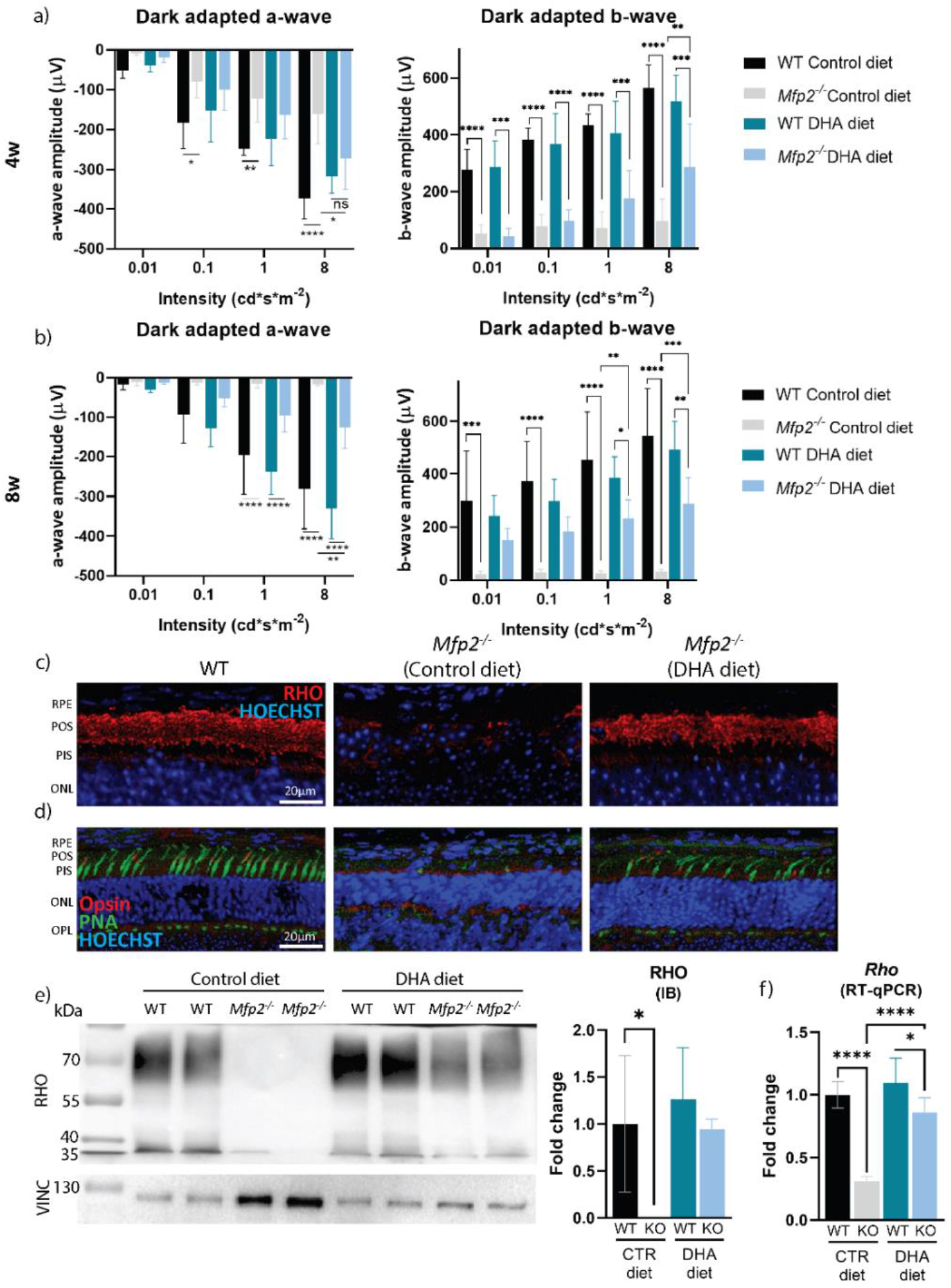
DHA supplementation improves photoreceptor functioning and morphology in Mfp2^−/−^ mice. **(a)** Dark adapted ERG responses from 4- and **(b)** 8-week-old mice. **(c)** Rod-specific stainings visualized with rhodopsin (red) and **(d)** cone-specific staining visualized with opsin (red) and peanut agglutinin lectin (PNA) (green) on 8-week-old retinal sections. **(e)** Immunoblotting (IB) and quantification of rhodopsin protein (8w). **(f)** RT-qPCR for rhodopsin mRNA levels (8w). N=4-8/group. Statistical difference based on multiple two-way ANOVA for the ERGs and one-way ANOVA for immunoblotting and RT-qPCR. Error bars indicate SD. RPE—retinal pigment epithelium; POS—photoreceptor outer segments; PIS—photoreceptor inner segments; ONL—outer nuclear layer; OPL—outer plexiform layer; RHO—rhodopsin; PNA—peanut agglutinin lectin; IB—immunoblotting; VINC— vinculin; CTR—control; ∗p < 0.05; ∗∗p < 0.01; ∗∗∗p < 0.001; ∗∗∗∗p < 0.0001.

To investigate the underlying reason for the declined rod photoreceptor responses at 8w, the localization and levels of the rod photoreceptor-specific marker, rhodopsin, were assessed in 8-week-old mice. In line with the H&E stainings (Figure 2b), IHC staining showed an impressive improvement in rod photoreceptor length in DHA supplemented *Mfp2^−/−^* mice compared to *Mfp2^−/−^*mice on control diet (Figure 3c). This was accompanied by normalisation of rhodopsin protein and even transcript levels (Figure 3e-f). Noteworthy, also cone photoreceptor morphology markedly improved in *Mfp2^−/−^* mice by DHA supplementation (8w) (Figure 3d). Overall, these findings reinforce previous reports that normal retinal DHA levels are essential for photoreceptor morphology and functioning^38–46^.

### DHA supplementation delays RPE dedifferentiation in *Mfp2^−/−^* mice

The RPE exerts an array of functions, with their main goal to maintain photoreceptor health. Accordingly, dysfunctions of the RPE cause degeneration of the retina. This was recently shown for mice lacking MFP2 specifically in the RPE (*Best1-Mfp2^−/−^* mice) (Kocherlakota et al. manuscript in revision). These mice presented with early onset RPE dedifferentiation similar to global *Mfp2^−/−^*mice. This consisted of loss of hexagonal shape, RPE depolarization, ablation of visual cycle proteins and RPE protrusions, causing secondary retinal degeneration at later ages (Kocherlakota et al. manuscript in revision). Therefore, it was of interest to investigate whether increased levels of DHA in the retina impacted on the RPE dedifferentiation process.

Strikingly, whereas the RPE of 8-week-old *Mfp2^−/−^* mice on control diet was strongly distorted, RPE cells maintained their hexagonal shape upon DHA supplementation, visualized with the tight junction marker zonula occludens protein 1 (ZO1) (Figure 4a). Furthermore, staining for ezrin, an apical marker that mislocalized to the basolateral side in *Mfp2^−/−^* mice on control diet, remained localized to the apical side in *Mfp2^−/−^* mice on DHA diet (Figure 4b). In addition, no RPE protrusions into the POS layer were seen (8w) (Figures 2b and 4b, white arrows). Moreover, transcript and protein levels of the crucial visual cycle protein 65 kDa retinoid isomerohydrolase (RPE65) were normal in the RPE of 4-week-old *Mfp2^−/−^* mice on DHA diet (Figure 4c, e). This was also the case for other visual cycle genes (Figure 4e). However, in contrast to the other RPE features, at 8w the visual cycle genes were suppressed by more than 60-90% compared to WT mice, despite DHA supplementation (Figure 4f). This coincided with a patchy signal for RPE65 in the RPE and severe loss of RPE65 protein, visualized with IHC and immunoblotting respectively (8w) (Figure 4b, d). Remarkably, the changes in visual cycle genes were not associated with alterations in the transcription factors *Sox9* or *Otx2*, which regulate their expression (Supplementary Figure 3)^47^. At 16w, the RPE dedifferentiated (including loss of hexagonal shape, RPE depolarization and RPE protrusions) despite continuous DHA supplementation (Figures 2c and 4a-b). These data indicate that DHA is important in maintaining RPE differentiation, but that other mechanisms are also involved.

**Figure 4.**
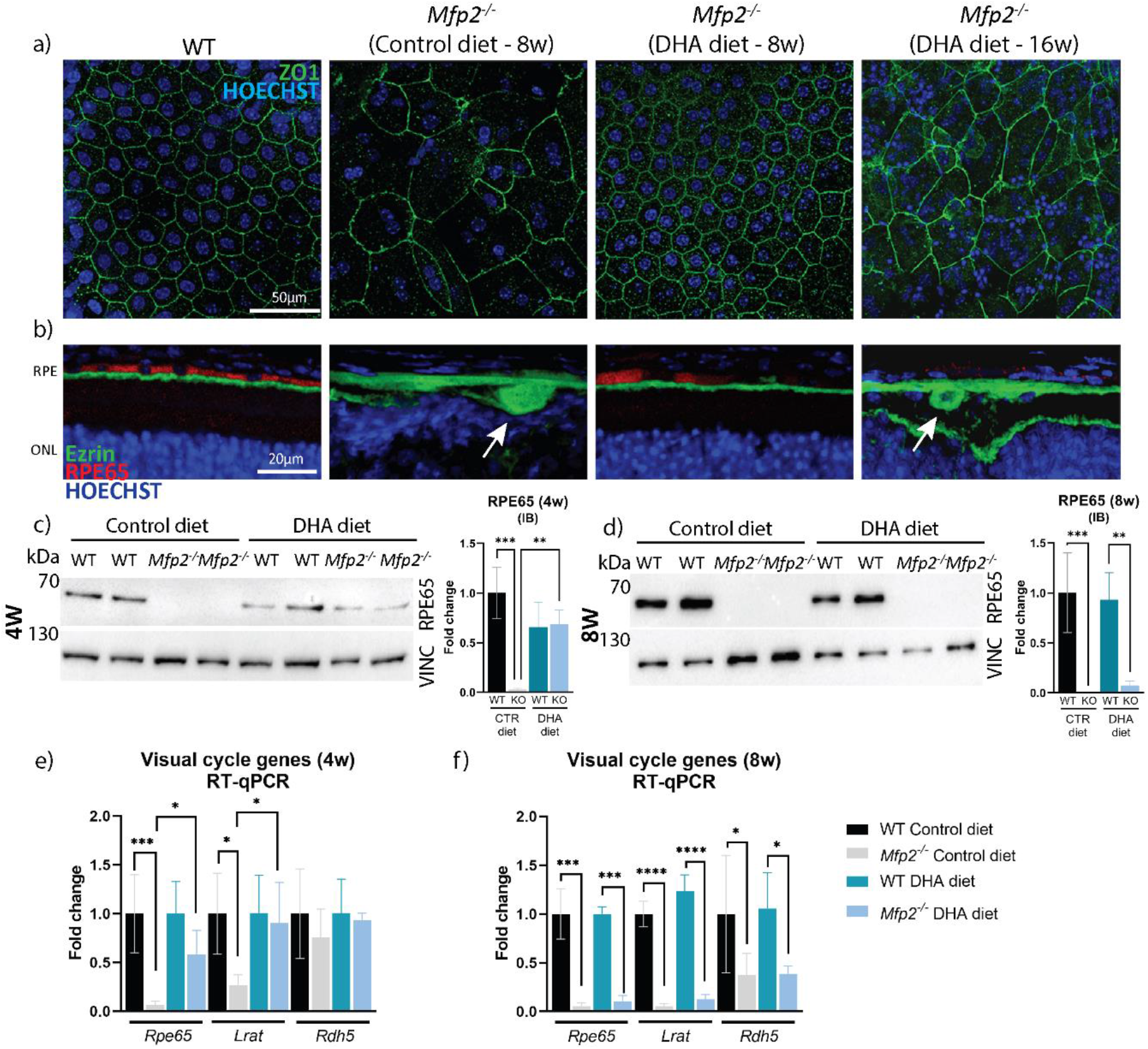
DHA supplementation delays RPE dedifferentiation in Mfp2^−/−^ mice. **(a)** ZO1 staining (green) on RPE wholemounts of 8- and 16-week-old mice. **(b)** IHC double staining for RPE65 (red) and ezrin (green) on 8- and 16-week-old mice. White arrows indicate RPE protrusions. **(c)** Immunoblotting (IB) for RPE65 in 4-week-old and **(d)** 8-week-old RPE samples. **(e)** RT-qPCR for the visual cycle genes at 4w and **(f)** 8w. N=4-7/group. Statistical difference based on multiple one-way ANOVA. Error bars indicate SD. LRAT—lecithin retinol acyltransferase; RDH5—retinol dehydrogenase 5; RPE—retinal pigment epithelium; ONL—outer nuclear layer; ZO1—zonula occludens-1; RPE65—65 kDa retinoid isomerohydrolase; IB—immunoblotting; VINC—vinculin; CTR—control; ∗p < 0.05; ∗∗p < 0.01; ∗∗∗p < 0.001; ∗∗∗∗p < 0.001.

### Supplementation of DHA affects the lysosomal functioning of the *Mfp2^−/−^* RPE

Another essential function of the RPE is the daily phagocytosis of damaged POS. Importantly, we recently demonstrated that loss of peroxisomal β-oxidation in the RPE impairs the digestive function of lysosomes, whereby both the degradation of POS phagosomes and of autophagic cargo was hampered. The prolonged presence of undigested POS caused overactivation of mammalian target of rapamycin (mTOR), which is known to trigger RPE dedifferentiation (Kocherlakota et al. manuscript in revision). Hence, it was of interest to evaluate if DHA supplementation influenced the lysosomal function of the *Mfp2^−/−^* RPE.

Characteristic of dysfunctional lysosomes in the RPE is the accumulation of rhodopsin-containing POS particles^35^, which we previously demonstrated to accrue in C57Bl6 mice (Kocherlakota et al. manuscript in revision). Similarly, more rhodopsin-containing POS particles were observed in the RPE of 4-week-old *Mfp2^−/−^* mice on control diet (Figure 5a). Intriguingly, this accumulation was not observed in the *Mfp2^−/−^* RPE supplemented with DHA at the same age, indicating the improved ability of the RPE to digest the POS (Figure 5a). Next, the status of mTOR was evaluated, by assessing levels of the phosphorylated form of its downstream target s6 (P-s6). In line with the notion that mTOR hyperactivation is caused by undigested POS, P-s6 levels were not upregulated in the *Mfp2^−/−^* RPE on DHA diet (Figure 5c). Finally, the lysosomal degradation of autophagic cargo was evaluated, by measuring levels of p62, a protein which binds and marks autophagic cargo for degradation. Indeed, while p62 vastly accumulated in the RPE of *Mfp2^−/−^* mice on control diet (4-fold), this did not occur in *Mfp2^−/−^* mice on DHA diet at 4w (Figure 5c). These findings indicate that DHA supplementation rescues the lysosomal functioning of the *Mfp2^−/−^* RPE at 4w. However, the rescue of lysosomal functioning was only temporary, as 8-week-old *Mfp2^−/−^* RPE cells supplemented with DHA diet showed POS accumulation, hyperactivated mTOR and higher p62 levels (Figure 5b, d). Of note, the number of rhodopsin-positive POS in the *Mfp2^−/−^* RPE on control diet normalizes to WT levels at 8w, most likely as there are no POS left to be digested as the retina is completely degenerated (Figure 5b).

**Figure 5.**
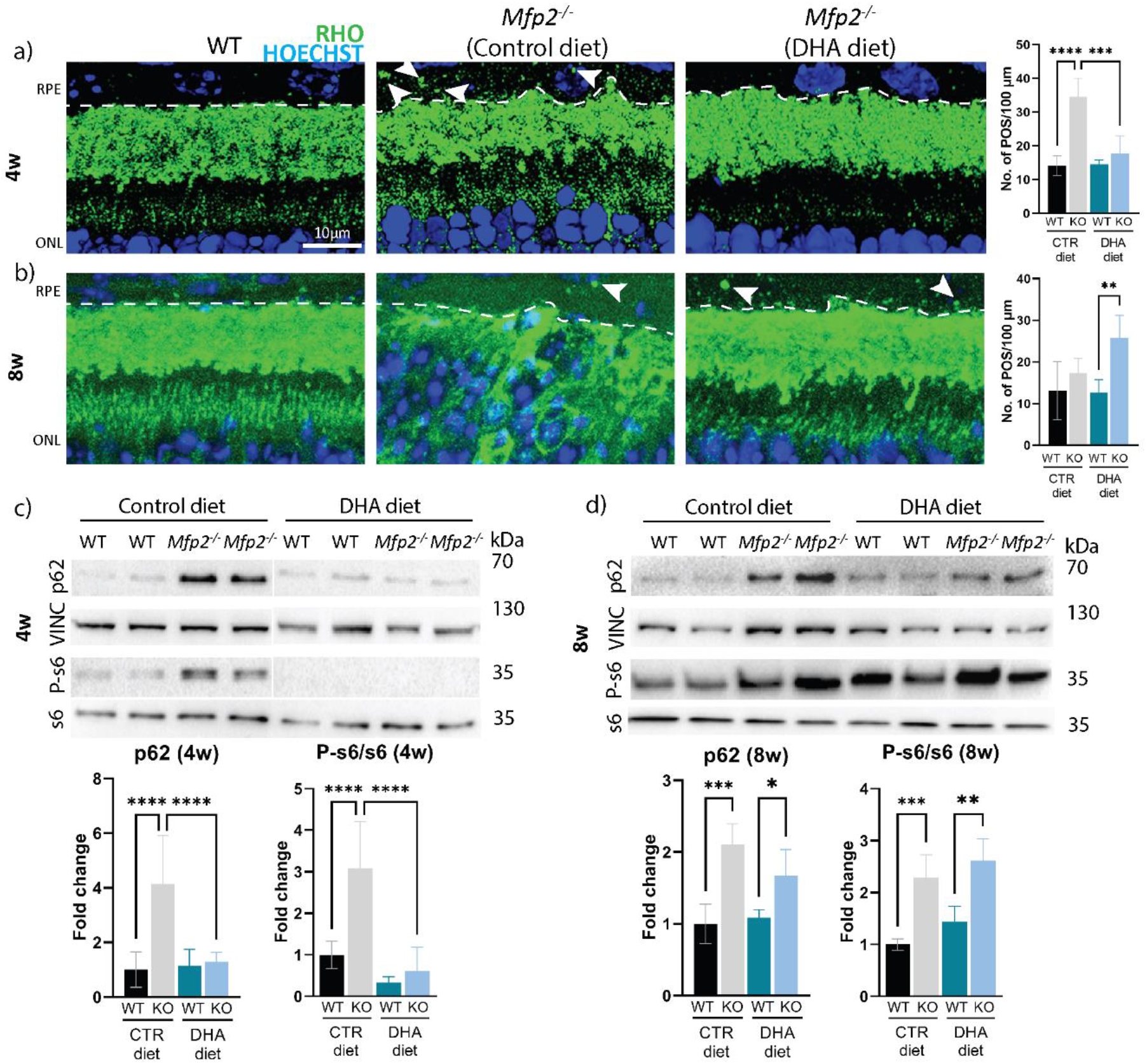
Improved RPE functioning in 4-week-old DHA-supplemented Mfp2^−/−^ mice. **(a)** IHC staining on 4-week-old and **(b)** 8-week-old mice for rhodopsin-containing POS particles (green)(illustrative instances are indicated with white arrows) in the RPE (delineated with dotted line). Quantifications of rhodopsin-positive POS per 100 µm are located on the right, per respective age. **(c)** Immunoblotting for p62, P-s6 and s6 on 4-week-old and **(d)** 8-week-old RPE samples. Vinculin was used as loading control. N=4-8/group. Statistical difference based on multiple one-way ANOVA. Error bars indicate SD. RPE—retinal pigment epithelium; ONL—outer nuclear layer; RHO—rhodopsin; P-s6—phosphorylated ribosomal protein s6; VINC—vinculin; CTR—control; ∗p < 0.05; ∗∗p < 0.01; ∗∗∗ - p < 0.001; ∗∗∗∗ - p < 0.0001.

Taken together, these results imply that the depletion of DHA in the *Mfp2^−/−^* retina on control diet contributed to the lysosomal dysfunction in the RPE at an early age. Furthermore, the temporary rescue of this endolysosomal system could play a role in the delay of RPE dedifferentiation onset in the DHA supplemented *Mfp2^−/−^* mice.

## Discussion

In this manuscript, we established that the early retinal phenotype of *Mfp2^−/−^* mice is inflicted by an impaired systemic supply of DHA. On the other hand, the retinopathy at later ages is most likely due to the inability of the RPE to handle the VLC-PUFA-containing POS, as a result of impaired peroxisomal β-oxidation. Altogether, this research provides a better understanding of the factors causing the retinopathy in peroxisome-deficient patients and offers insight into the distinct roles of DHA in the neural retina and RPE cells.

Firstly, in this manuscript we confirm that adequate levels of retinal DHA are required for normal postnatal photoreceptor development and function. Furthermore, the rescue of the early *Mfp2^−/−^* photoreceptor phenotype via increasing the systemic supply of DHA, highlights the involvement of DHA in the early retinal degeneration in *Mfp2^−/−^*mice. These conclusions are in line with other mouse models with a genetic defect in the acquisition of retinal DHA, showing a similar retinal phenotype^4^. However, the molecular details by which DHA influences photoreceptor homeostasis is still not fully understood. Although the role of DHA in photoreceptor biogenesis and phototransduction has been well studied, its role in oxidative stress is controversial ^2,4,48,49^. On the one hand, it was shown that under uncompensated oxidative stress, RPE cells convert DHA to the protective lipid mediators NPD1, resolvins, protectins and maresins^50^. On the other hand, the high content of double bonds in this PUFA predisposes to lipid peroxidation, which can be deleterious for photoreceptors^51–54^. It was suggested that the impact of DHA may depend on the circumstances such as the level of oxidative stress^55^. Interestingly, targeted metabolome analysis on *Mfp2^−/−^* RPE revealed that levels of redox metabolites were unchanged at 3w (Kocherlakota et al. manuscript in revision). In agreement, despite several attempts, we failed to reliably measure the protective lipid mediators in (DHA supplemented) *Mfp2^−/−^* mice. However, we cannot exclude that, upon DHA supplementation, more NPD1 was generated in the *Mfp2^−/−^*retina, thereby contributing to the delay of the retinal degeneration. To investigate a potential protective role of NPD1, *Mfp2^−/−^* mice could be supplemented with this mediator.

Lipidome analysis on the neural retina of DHA supplemented *Mfp2^−/−^* mice, revealed a striking accumulation of VLC-PUFA-containing phospholipid species already at the age of 4w. Interestingly, this accumulation was more pronounced in the DHA supplemented *Mfp2^−/−^* mice, compared to *Mfp2^−/−^* mice on control diet. This confirms our previous findings that the vast accretion of VLC-PUFAs does not initiate the early retinopathy in *Mfp2^−/−^* mice^25^. However, the inability to process the VLC-PUFAs due to dysfunctional peroxisomal β-oxidation in the RPE, could cause the VLC-PUFAs to reach a toxic threshold, thereby contributing to the destruction of the retina at 16w.

Remarkably, VLC-PUFAs were differentially distributed in the *Mfp2^−/−^* retina upon DHA diet. PC-containing VLC-PUFAs (≥C38) were further increased by the diet in *Mfp2^−/−^* retina, whereas TG species containing these VLC-PUFAs were partially reduced, coinciding with lowered lipid droplet accumulation in both the RPE and neural retina. So far, the mechanism by which DHA affects the storage of fatty acids remains unknown. It would be interesting to track the fate of PUFAs and gain insight into the composition of the lipid droplets in the (DHA supplemented) *Mfp2^−/−^*retinas, via applying the recently developed techniques i) MALDI-MSI^56–59^, ii) colocalization studies of injected photoreactive lipid probes^60^ and lipid droplets visualized with superresolution microscopy, or iii) *in vivo* administration of radio-labelled DHA combined with spatial lipidomics^61^.

The most intriguing and unexpected finding was that DHA temporarily improved RPE homeostasis. This seems indeed in contradiction with the cell-autonomous role of MFP2 in the RPE, shown by using *Best1-Mfp2^-/-^* mice (Kocherlakota et al. manuscript in revision).

The normal lysosomal function and reduced lipid droplet accumulation in the DHA supplemented *Mfp2^−/−^* RPE at 4w, despite a 20-200 fold accumulation of VLC-PUFA-containing lipid species in their neural retina, indicated that the *Mfp2^−/−^* RPE supplemented with DHA are somehow able to handle the VLC-PUFA-containing POS. Interestingly, POS accumulation in the RPE was also observed in other DHA-deficient mouse models, e.g. *AdipoR1^−/−^*^62^ and *Mfsd2a^−/−^*mice^63^, suggesting that DHA might play a role in the phagocytosis process. Perhaps DHA influences the membrane characteristics, thereby affecting the breakdown of POS phagosomes. However, despite continuous DHA supplementation, the lysosomal function subsequently declined. To gain insight into the temporary rescue of lysosomal functioning, it will be important to determine the sequence of impaired lysosomal function versus accumulation of lipids in the *Mfp2^−/−^* RPE. *In vitro* studies supplementing normal POS versus DHA-deprived POS to healthy and *Mfp2^−/−^* RPE cells could shed some light on the mechanisms of RPE disruptions and the role of DHA in POS phagocytosis. Nevertheless, it remains unsolved how the DHA supplemented *Mfp2^−/−^* RPE is able to handle the VLC-PUFAs present in the POS.

The transient rescue of the visual cycle genes was remarkable and provided several essential insights. Firstly, the normal levels of visual cycle genes at 4w and suppression at 8w in DHA supplemented *Mfp2^−/−^* mice correlated with the respectively normal and impaired scotopic a-wave responses at these ages. Since the visual cycle genes play a crucial role in the phototransduction, it is plausible that alterations in their levels impaired the rod photoreceptor responses of the DHA supplemented *Mfp2^−/−^* mice at 8w. Secondly, both RPE65 depletion^64^ and dedifferentiation of the RPE are known causes for retinal degeneration^65^, implying that these factors contributed to the retinopathy at 16w in the DHA supplemented *Mfp2^−/−^*mice. However, it remains to be determined what drives the reduction of the visual cycle genes. Furthermore, it remains unclear if the reduction in visual cycle genes is either the first sign of RPE dedifferentiation or an independent event.

The short-term rescue of the RPE phenotype in *Mfp2^−/−^*mice upon DHA supplementation is supported by other mouse models with a genetic defect in retinal PUFA acquisition, in which several RPE abnormalities were reported^4^. However, it remains to be determined whether the RPE phenotype is due to a primary deficiency of DHA levels in the RPE or a secondary effect of neural retina degeneration. Interestingly, comparison of the retinal phenotype of *Crx-Mfp2^−/−^* ^25^ and *Best1-Mfp2^−/−^* mice (Kocherlakota et al. manuscript in revision), indicated that loss of MFP2 from the RPE, and not from photoreceptors, is detrimental for the retina. Taken together, it seems worthwhile to further explore the role of DHA in the RPE.

What are the implications of this research for therapeutic approaches for peroxisome-deficient patients? Major drawbacks with regard to retinal research in peroxisomal disorders is translatability of the findings from the mouse models to patients, due to i) the lack of reports on the histopathological changes and lipid content of the retina of peroxisome-deficient patients, ii) the promising, but inconclusive results of the clinical trials with DHA supplementation to peroxisome-deficient patients^66,67^, and iii) the differences in cone density between human and mice retinas. This hinders to draw conclusions on the causative factors underlying the retinopathy in peroxisome-deficient patients, and thus the ability to recommend a treatment option. Nevertheless, the present data imply that sole DHA supplementation will not be able to prevent the retinopathy in patients with peroxisome deficiencies. Therefore, a combination approach of DHA supplementation to normalize the systemic DHA supply, together with local viral delivery of the missing gene (as already reported for *Pex1*-mutant mice^68^) in order to process the (VLC-)PUFAs in the RPE and neural retina, seems plausible.

## Funding

This research was funded by Belgian Fund for Research in Ophthalmology, by the KU Leuven (C14/18/088) and by the Research Foundation—Flanders (FWO G0A8619N). The Leica SP8x confocal microscope was provided by InfraMouse (KU Leuven-VIB) through a Hercules type 3 project (ZW09-03).

## Commercial relationship disclosure

The authors declare no competing interests.

## Supplementary figures

**Figure S1.**
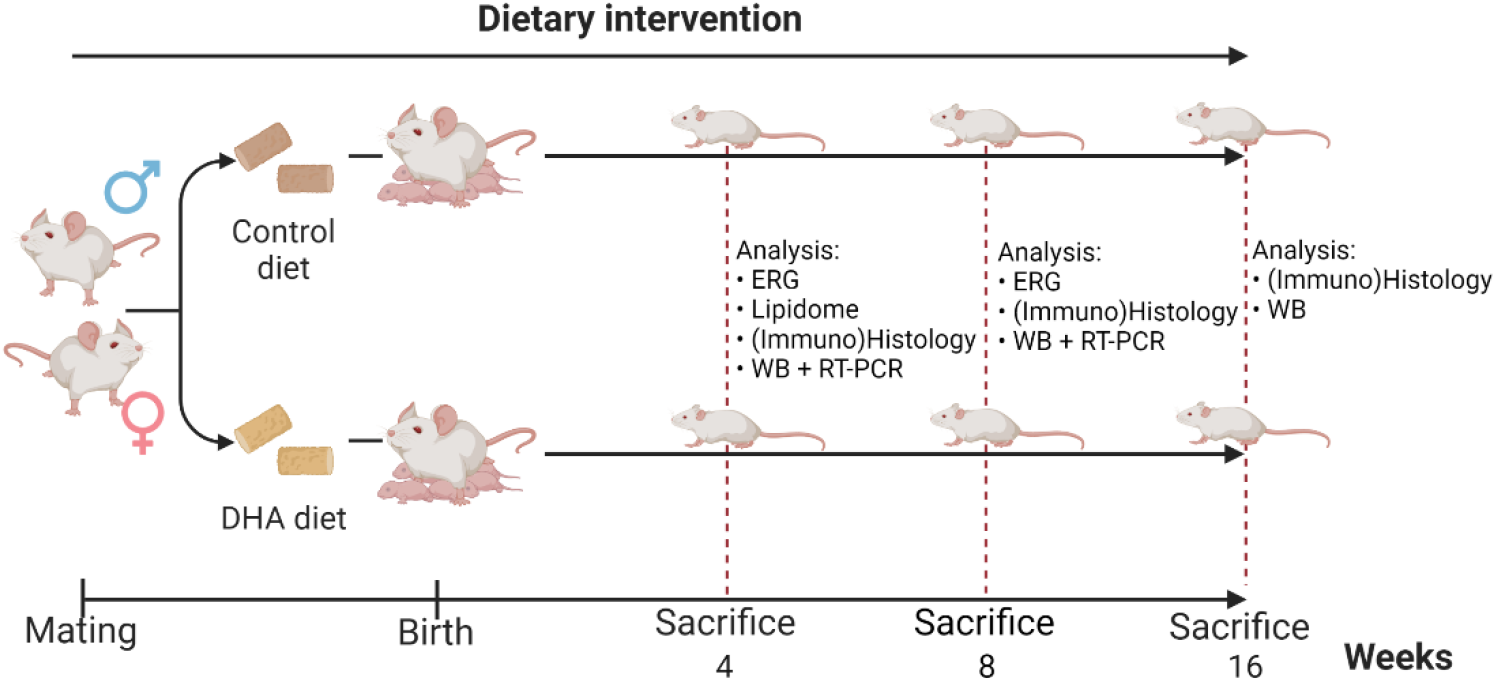
DHA supplementation to Mfp2^−/−^ mice. At the moment of mating, breeding pairs were put on either a control or DHA diet. Consequently, the Mfp2^−/−^ pups and controls already receive the diet during the gestation and lactation period. After weaning (±P21), offspring received the diet until sacrifice. Figure created with Biorender.com.

**Figure S2.**
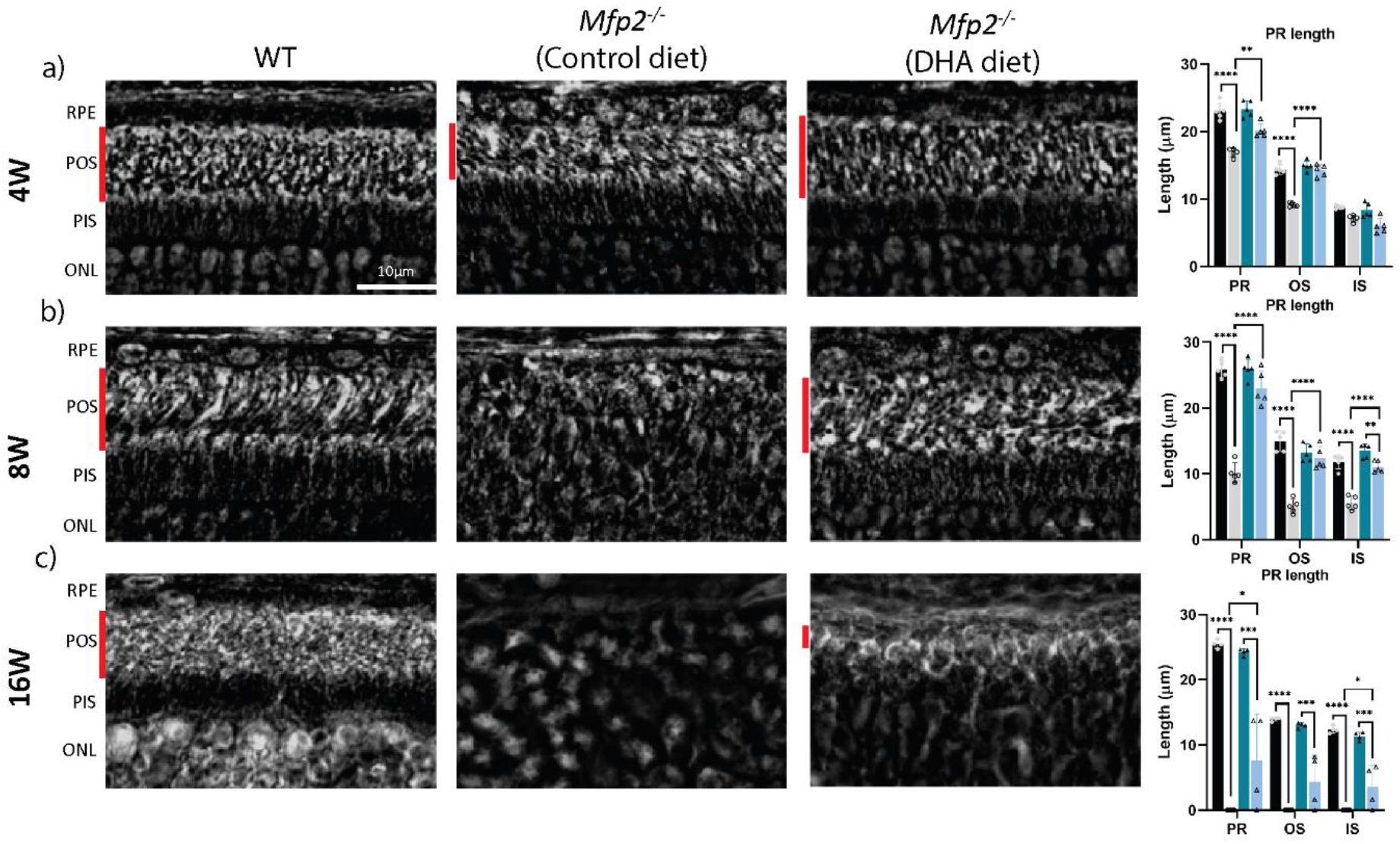
Delayed outer segment shortening in Mfp2^−/−^ mice on DHA diet. Measurement of photoreceptor layers on phase contrast microscopy images in 4-**(a)**, 8-**(b),** and 16-week-old **(c)** mice. Red bars indicate POS length. N=4-5/group. Statistical difference based on multiple one-way ANOVA. Error bars indicate SD. RPE— retinal pigment epithelium; POS—photoreceptor outer segments; PIS—photoreceptor inner segments; ONL— outer nuclear layer; PR—photoreceptor. * p < 0.05, ** p < 0.01, *** p < 0.001, **** p < 0.0001.

**Figure S3.**
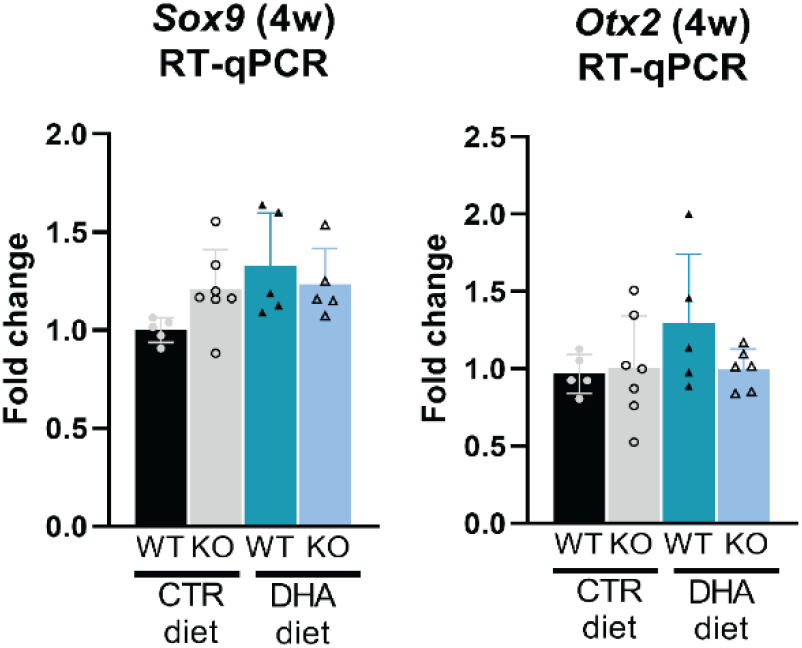
RT-qPCR for transcription factors regulating visual cycle protein expression (4w). N=5-8/group. Statistical difference based on multiple one-way ANOVA. Error bars indicate SD. Otx2—orthodenticle homeobox 2; Sox9—SRY-box transcription factor 9.

## Supplementary tables

**Supplementary Table 1.**
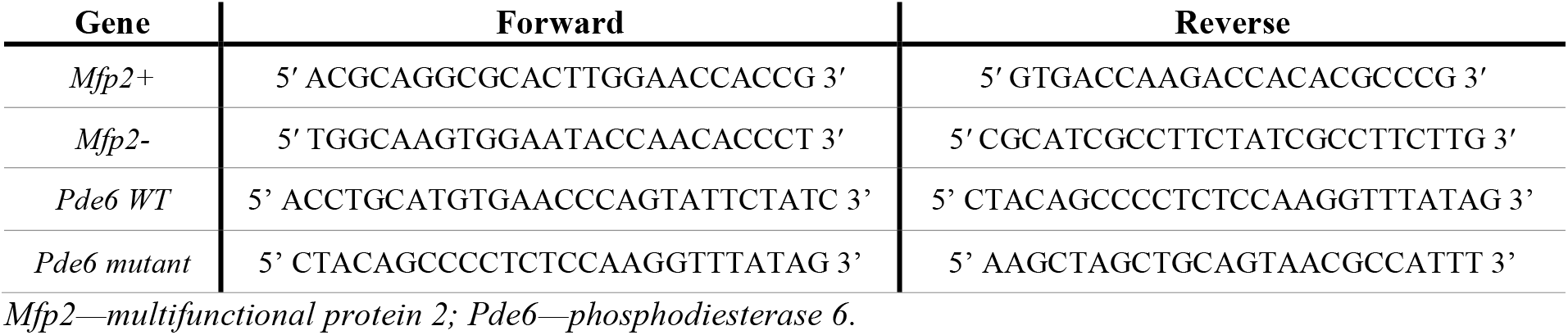
List of primers used for PCR genotyping.

**Supplementary Table 2.**
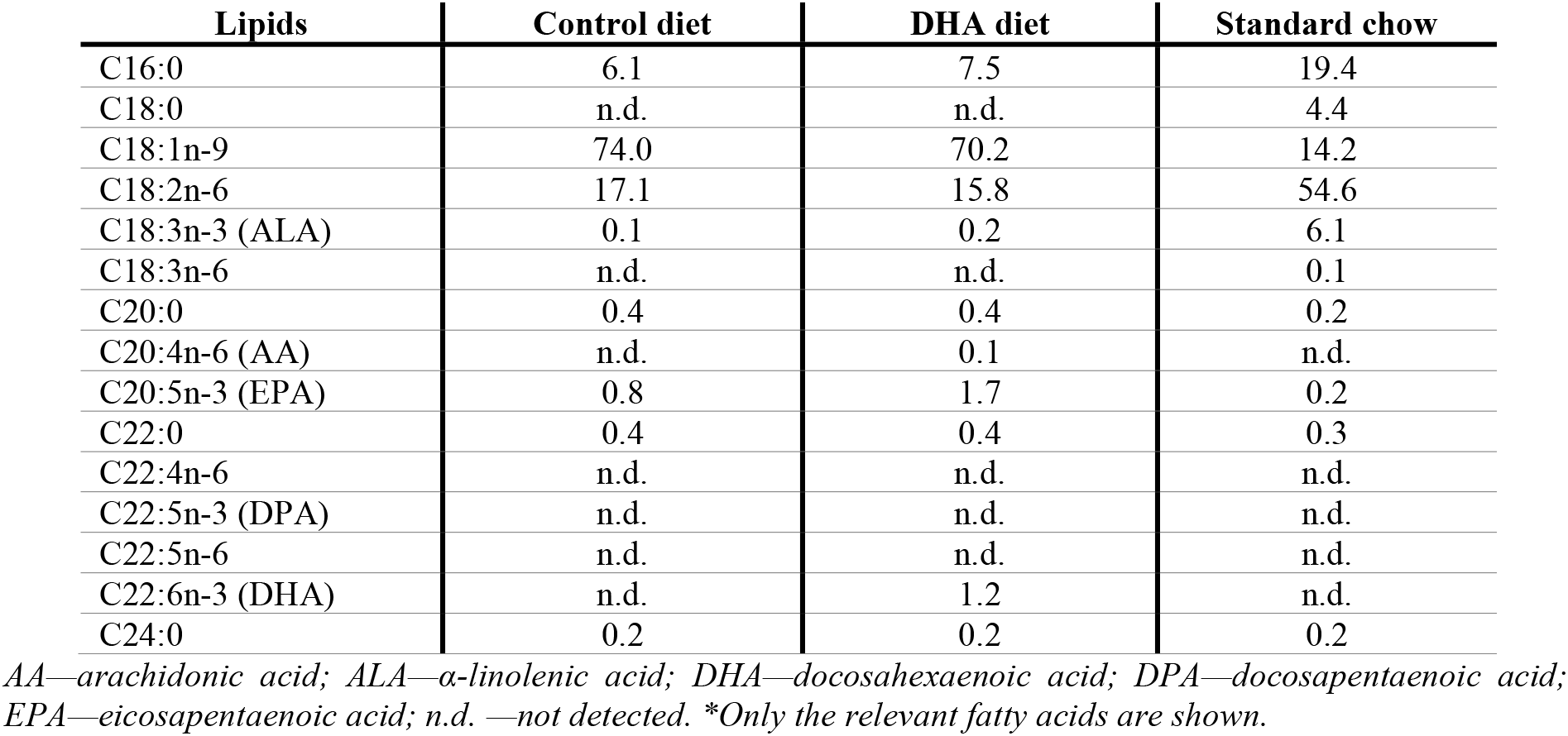
Fatty acid composition of control, DHA and standard diet ((%w/w of total fatty acids)*

| Lipids | Control diet | DHA diet | Standard chow |
| --- | --- | --- | --- |
| C16:0 | 6.1 | 7.5 | 19.4 |
| C18:0 | n.d. | n.d. | 4.4 |
| C18:1n-9 | 74.0 | 70.2 | 14.2 |
| C18:2n-6 | 17.1 | 15.8 | 54.6 |
| C18:3n-3 (ALA) | 0.1 | 0.2 | 6.1 |
| C18:3n-6 | n.d. | n.d. | 0.1 |
| C20:0 | 0.4 | 0.4 | 0.2 |
| C20:4n-6 (AA) | n.d. | 0.1 | n.d. |
| C20:5n-3 (EPA) | 0.8 | 1.7 | 0.2 |
| C22:0 | 0.4 | 0.4 | 0.3 |
| C22:4n-6 | n.d. | n.d. | n.d. |
| C22:5n-3 (DPA) | n.d. | n.d. | n.d. |
| C22:5n-6 | n.d. | n.d. | n.d. |
| C22:6n-3 (DHA) | n.d. | 1.2 | n.d. |
| C24:0 | 0.2 | 0.2 | 0.2 |

AA—arachidonic acid; ALA— α-linolenic acid; DHA—docosahexaenoic acid; DPA—docosapentaenoic acid; EPA—eicosapentaenoic acid; n.d. —not detected. \*Only the relevant fatty acids are shown.

**Supplementary Table 3.**
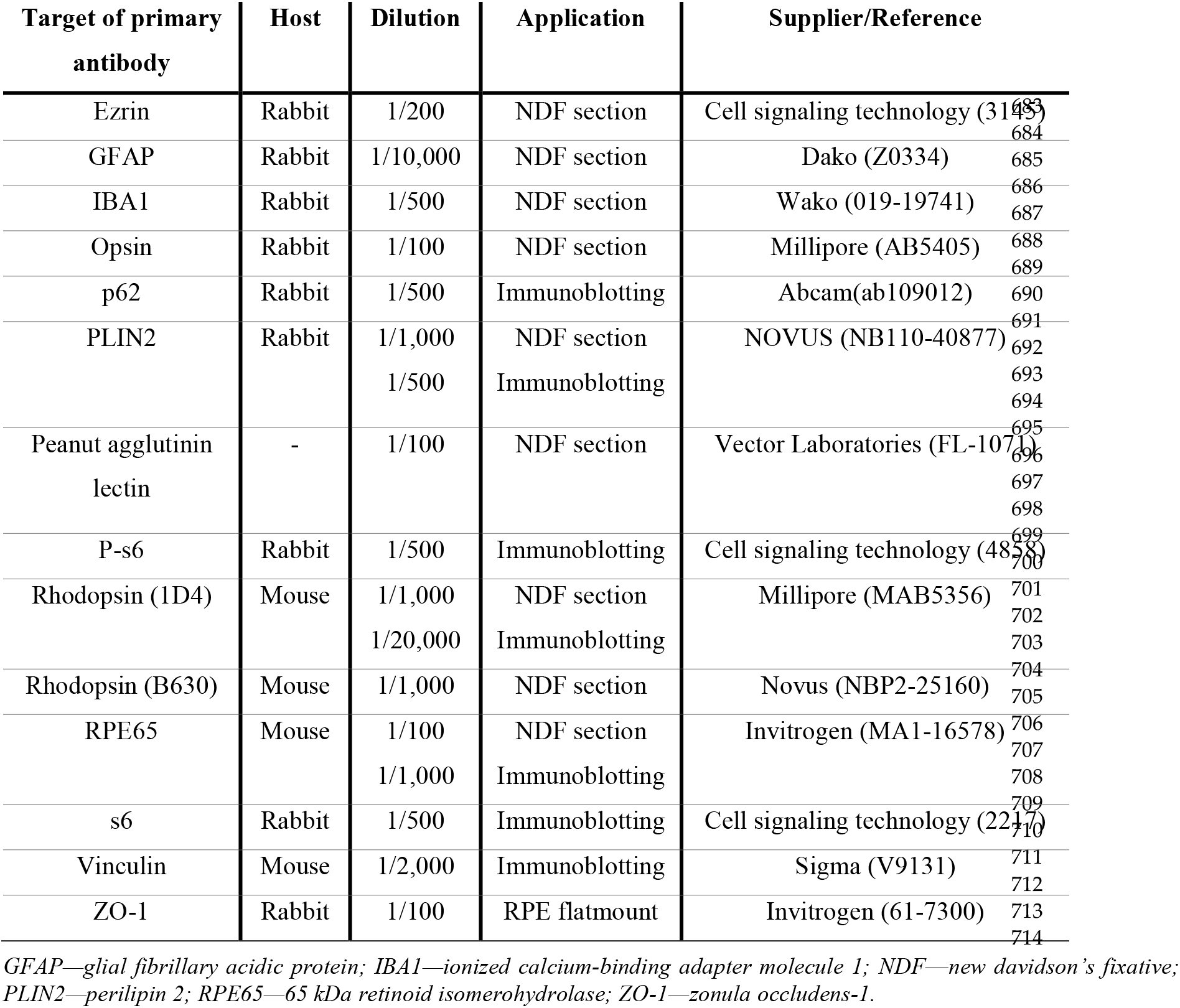
List of used primary antibodies for immunohistochemical stainings and immunoblotting.

**Supplementary Table 4.**
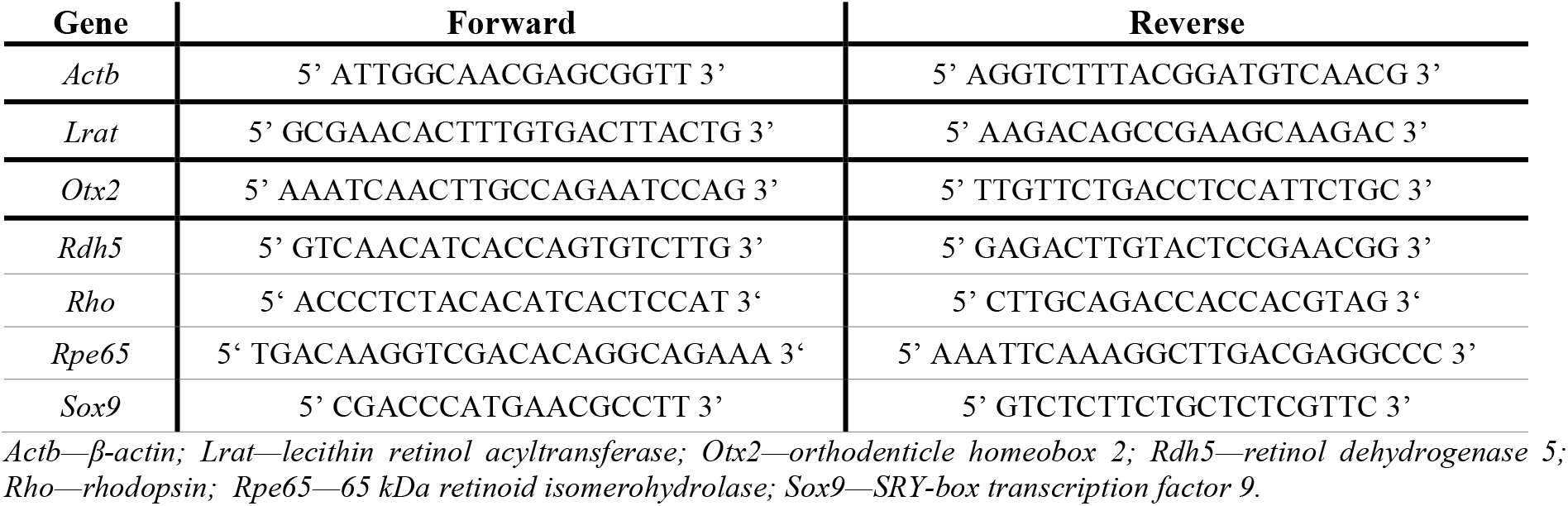
List of primers used for RT-qPCR.

## Acknowledgements

The authors thank E. Nefyodova, B. Das, A. Carton and A. Manderveld for the excellent technical assistance. This research was funded by the KU Leuven (C14/18/088) and by the Research Foundation—Flanders (FWO G0A8619N). The Leica SP8x confocal microscope was provided by InfraMouse (KU Leuven-VIB) through a Hercules type 3 project (ZW09-03).

## Notes

### Competing Interest Statement

The authors have declared no competing interest.

